# MAFA Phosphorylation Controls Beta-Cell Identity and Sex-Specific Pancreatic Disease Outcomes

**DOI:** 10.1101/2025.06.11.659123

**Authors:** Moustapha Ka, Sabine Druillennec, Sara Chabi, Céline Roullé, Dave YX Ng, Christine Péchoux, Cédric Messaoudi, Christine Walczak, Marie-Noëlle Soler, Laetitia Besse, Laure Lecoin, Sandra Majo, Gérard Gradwohl, Alain Eychène, Benoit Terris, Celio Pouponnot, Bertrand Duvillié

## Abstract

*Mafa* is a critical transcription factor in pancreatic beta-cell biology, orchestrating insulin expression in response to glucose elevations. As a member of the large MAF protein family, MAFA’s stability and activity are intricately regulated by GSK3-mediated phosphorylation. To decipher the functional roles of these phosphorylations, we engineered knock-in mice (*Mafa^4A^*^/+^) in which MAFA is rendered non-phosphorylatable.

In all *Mafa^4A^*^/+^ animals, MAFA stability was markedly enhanced. Under high-fat diet (HFD) conditions, *Mafa^4A^*^/+^ males rapidly developed glucose intolerance, which was attributed to impaired glucose-stimulated insulin secretion. Bulk RNA sequencing revealed disrupted beta-cell identity, characterized by increased expression of MODY-associated genes and a delta-cell signature, suggesting beta-to-delta cell reprogramming, a hypothesis supported by lineage-tracing experiments.

Conversely, *Mafa^4A^*^/+^ females exhibited hypoglycemia and, with age, developed pronounced inflammatory cystic ducts including mucinous cystic neoplasms (MCNs). Strikingly, MAFA protein was also detected in MCN biopsies from female patients, linking our findings to human pathology.

Our results unveil a sex-biased impact of GSK3-mediated MAFA phosphorylation. The male phenotype closely parallels the MODY-like diabetes observed in patients with MAFA S64F mutations, implicating defective phosphorylation in disease etiology. The emergence of MCNs in female mice suggests a novel role for MAFA stability or mutations in the pathogenesis of these enigmatic neoplasms, providing a fresh molecular hypothesis with clinical relevance.

## Introduction

*Mafa* (musculoaponeurotic fibrosarcoma oncogene homolog A) is a transcription factor which belongs to the large *Maf* transcription factor family (1). Large *Maf* are oncogenes involved in developmental, metabolic, and tumor processes (2,3). Their transforming and differentiating activity are regulated by GSK3-mediated phosphorylation which depends on a priming phosphorylation (4,5). Mafa is first phosphorylated by an unknown priming kinase on serine 65 (S65) allowing the subsequent phosphorylations by GSK3 on 4 serine and threonine residues (S61, T57, T53, and S49). This induces ubiquitination and degradation of MAFA, but paradoxically, it also enhances its transcriptional activity (6).

In the pancreas, *Mafa* is a key transcription factor of beta cell identity where it plays a key role in their maturation and pathophysiology. MAFA was first shown to bind the Maf recognition element (MARE) within the RIPE3b/C1-C2 sequence of the insulin promoter, thereby regulating insulin expression in response to glucose (7–9). In mice, *Mafa* expression begins at E13.5 in developing beta cells and is maintained in adult mature beta cells (10). *Mafa* knock-out mice develop impaired glucose-stimulated insulin secretion (GSIS), leading to diabetes at 50 weeks (11,12). In the absence of *Mafa*, beta cells display decreased expression of numerous transcription factors that are crucial determinants of beta cell identity and function. This includes the expression of *Ins1*, *Ins2*, *Pdx1*, *NeuroD1*, and *Slc2a2* (13). Other MAFA target genes have also been uncovered (14). In addition, *Mafa* is important for beta cell maturation. Indeed, neonatal rat islets are immature and express low levels of *Mafa*. Overexpression of *Mafa* in these neonatal islets enhances GSIS, confirming their progression to a more mature state (15).

In humans, several *Mafa* mutations have also been associated with diabetes. Indeed, *Mafa* expression is decreased in patients with type 2 diabetes (T2D), and a *Mafa* polymorphism is also associated with type 1 diabetes (T1D) (16). More recently, a MAFA S64F mutation was found in two independent families. Patients carrying the S64F mutation develop either diabetes, predominantly in males, or insulinomatosis, more frequently in females (17). The male phenotype has also been mostly replicated in mice carrying the same S64F mutation (18). According to the proximity to S65, the S64F mutation has been shown to increase MAFA stability and decrease GSK3-mediated phosphorylation. Based on these data, one hypothesis is that the S64F mutation might prevent the priming kinase from phosphorylating S65 and subsequent GSK3-mediated phosphorylation.

To investigate the specific role of GSK3-mediated phosphorylation of MAFA, we generated a knock-in *Mafa^4A^* mutant mice in which all four serine/threonine residues are mutated to alanine. This mutated form of MAFA can no longer be phosphorylated by GSK3 and degraded by the proteasome. As a result, it increases its accumulation and alters its transcriptional capacity. Homozygous *Maf*a^4A/4A^ mice die at birth of high incidence of breath holding fatal apneas (19). We made use of the heterozygous *Mafa*^4A/+^ knock-in mice to study the role of GSK3-mediated MAFA phosphorylation in pancreatic beta cells. Our KI model allows us to understand whether the biological effect of S64F mutation is due to the prevention of GSK3-mediated phosphorylations.

## METHODS

### Animals

The *Mafa^flox4A^*^/+^mice were generated as described in (19) on the 129 Sv genetic background. These floxed mice were first crossed with the pPGK-Cre line to obtain ubiquitously defloxed *Mafa*^4A/+^ mice. These knock-in mice express the non-phosphorylatable form of MAFA (*Mafa*^4A/+^). Of note, homozygous *Mafa^4A/4A^* mice die at birth from respiratory distress (19). For this reason, we focused our study on heterozygous mice. For the lineage tracing experiments, we used pIns1-Cre^ERT2^ (20) generated at the ICS (Illkirch, France), mT/mG and *Mafa*^4Afl/+^ mice on the C57/Bl6J background. Mice were group-housed with free access to food and water in controlled conditions (temperature 21°C, humidity 40-50%) and exposed to conventional 12 h light/dark cycle. For high fat diet experiments, the mice were exposed to 60 kcal% fat (Research Diet, Lynge, Denmark).

### Study Approvements

All the experimental procedures were performed in accordance with the European recommendations and the Local Ethics Committee (Authorization DAP 2018-015).

### Glucose and insulin tolerance tests

The mice were first fasted for 16 hours for the glucose tolerance test (GTT) and 6 hours for the insulin tolerance test (ITT). After being weighed individually, 2 g of glucose (Lavoisier, Paris, France) were injected intraperitoneally per kg per mice for the GTT. On the other hand, 0.75U of insulin per kg was injected for the ITT. Blood glucose was measured using a glucometer (Accucheck Performa, Roche, Boulogne Billancourt, France) at 0, 15, 30, 60, 90 and 120 min after the glucose injection for the GTT, and at 0, 15, 30 and 60 min after the insulin injection for ITT.

### Pancreatic Islet Isolation

The islets of Langerhans were prepared as described in (21) with some modifications. 2 ml of collagenase 1mg/ml (Sigma-Aldrich C7657, Merck, France) associated with DNase 1 mg/ml (DNase 1 Roche 10104159001, Merck, France) was injected into the pancreas through the common bile duct. The pancreas was then digested in a water bath (37°C) for 22 min in 3 ml of additional collagenase. Digestion was stopped by the addition of 25 ml of a solution of HBSS Ca^2+^ and Mg^2+^ (Gibco 14065-072, Thermo Fisher, France) containing 10% fetal calf serum. The mixture was next centrifuged for 30 seconds at 290 g. The supernatant was removed, and the pellet was washed twice in HBSS containing Ca^2+^, Mg^2+^ with 10% serum. The pellet was resuspended in 10 ml of RPMI (Gibco 1640, Thermo Fisher, France) and vortexed. 10 ml of histopaque (density of 1.077 g/ml, Sigma Aldrich 1077, Merck, France) was added gently to obtain two distinct phases. The mixture was centrifuged at 922 g for 27 min at room temperature. After centrifugation, the islets were located at the interface between the histopaque and RPMI. They were collected and washed 3 times with 10% HBSS serum.

### Insulin secretion test

Islets were cultured overnight in 2 ml RPMI 1640 (Thermo Fisher) with 100 units/ml penicillin, 100 ug/ml streptomycin (Gibco) and BSA (5 g/l). Batches of 5 islets were taken to perform the secretion test. The islets were washed in 1 ml of 0.5% (w/vol) bovine serum albumin in Krebs-Ringer Hepes-Buffered saline (Thermo Fischer) with 2.8 mM glucose at 37°C in 5% CO2. Next, the islets were transferred sequentially in 1 ml of 0.5% (w/vol) bovine serum albumin in Krebs-Ringer Hepes-Buffered saline with 2.8 mM glucose for one hour, in 16.7 mM glucose for one hour and finally in 50mM KCl for 1 hour. After the secretion assay, the islets were digested in a solution of acid (1.5% HCl in 70% EtOH) to extract the insulin content. At all steps, the media were collected, and insulin was quantified by ELISA (Millipore, EZRMI-13K, Sigma).

### Immunofluorescence and immunohistochemistry

Tissues were fixed in 10% formalin and embedded in paraffin. Sections of 6 µm were collected and processed for immunofluorescence (IF) as described in (22). More precisely, slices were deparaffinized, rehydrated and microwaved for 4 min at 850 W and for 12 min at 145 W in Citra plus antigen retrieval buffer (Biogenex laboratories, Fremont, USA). Blocking and permeabilization were performed by incubating the slides in TBS containing 3% BSA and 0.3% Triton for 30 min. Primary and secondary antibodies used during all experiments are listed in Table 1 and 2 respectively. Bright-field and fluorescent images were acquired on a widefield AxioImager M2 upright microscope (Carl Zeiss, Germany) coupled with an AxioCam 506 color camera (Carl Zeiss, Germany). DAPI (ex BP 387/11; em BP 447/60) GFP (ex BP 470/40 ; em 525/50) Cy3 (ex BP 550/25 ; em 605/70) and TxRed (ex BP 500/20 ; em BP 535/30) were detected using bloc filters. The system is driven by Zen blue software. For confocal microscopy, images were acquired with a Leica SP8X inverted confocal laser scanning microscope (CLSM), equipped with a 63x Oil immersion objective (NA = 1.4). Sequential excitation mode (488 nm and 640 nm obtained with a white light laser (WLL)) was used to collect images on GaAsP Hybrid photon detectors. Emission was detected following excitation at 488 nm, with a detection range of 500–570 nm, and following excitation at 640 nm, with a detection range of 660–760 nm. The whole system was driven by the LAS X software (Leica).

### Alcian blue and HES staining

The staining of mucin uses the Wagner and Shapiro method (1957) (23). It employs the Alcian blue dye. The sections are deparaffinized and rehydrated with a decreasing alcohol series. Next, the sections are stained with Alcian blue at pH 2.6 for 20 minutes. After staining, the samples are rinsed in running water, then the nuclei are stained with nuclear red for 10 minutes and rinsed again in water. The sections are dehydrated with an increasing alcohol series, then incubated in xylene and mounted.

For Hematoxylin and eosin staining (HES), the sections are deparaffinized and rehydrated with a decreasing alcohol series and water. Next, the sections are stained with a hematoxylin solution (MM, France) for 1 minute and rinsed in water for a few minutes. Then, they are stained with 0.5% eosin (MM, France) and rinsed in water. An alcoholic saffron solution is used to stain the connective tissues for 5 minutes. Finally, the samples are dehydrated with absolute ethanol and then xylene, and mounted with Entellan.

### Transmission Electron Microscopy

Samples were fixed with 2% glutaraldehyde in 0.1 M Na cacodylate buffer pH 7.2, and then contrasted with Oolong Tea Extract (OTE) 0.2% in cacodylate buffer, postfixed with 1% osmium tetroxide containing 1.5% potassium cyanoferrate, gradually dehydrated in ethanol (30% to 100%) and substituted gradually in mix of ethanol-epon and embedded in Epon. Thin sections (70 nm) were collected onto 200 mesh copper grids, and counterstained with lead citrate.

Grids were examined with Hitachi HT7700 electron microscope operated at 80kV (Milexia – France), and images were acquired with a charge-coupled device camera (AMT), on MIMA2 facility - https://doi.org/10.15454/1.5572348210007727E12.

### Quantification

For the adult pancreases, quantification of insulin- and glucagon-staining were performed on five equally separated sections. In each section, the beta-cell, alpha-cell and pancreatic areas were determined using the NIH Image J (NIH, Washington, USA) software. The percentage area of beta-cells and alpha-cells in each pancreatic section was determined by dividing the area of all insulin-positive cells or glucagon-positive cells respectively by the total surface area of the section. The beta-cell mass and alpha-cell mass were calculated by multiplying the pancreas weight by the percentage area of the beta-cells and alpha-cells respectively. At E18.5, the entire pancreases were sectioned, and one of the two slides were analyzed. To quantify the absolute surface of insulin, somatostatin and DAPI, one out of three sections per slide were processed for IF. After digitalization, the surface of somatostatin, insulin and DAPI staining were measured for each section using Image J software. To measure the proliferation of beta-cells, we counted the frequency of nuclei among 1,000 insulin-positive cells. At least three rudiments per condition were analyzed.

The segmentation of secretion vesicles on transmission electron microscopy images was performed using Cellpose (24,25). The model cyto was refined using the human in the loop approach with a total of 21 images with vesicles at different stages from all conditions of the experiences. To analyze the full dataset of 288 images, the MIC-MAQ software (https://github.com/MultimodalImagingCenter/MIC-MAQ) was used to manage image segmentation with the trained model, export this segmentation as region of interest and perform quantifications. The distance to membrane was performed using ImageJ. A mask of each cell was given manually, the MorpholibJ plugin (26) was then used to compute an euclidian distance map were each pixel corresponds to the distance to the nearest membrane, and the minimum value is measured for each vesicle using the region of interest saved in the prior segmentation step.

### Statistics

GraphPad Prism 10.2.3 was used for all statistical analyses. All statistical methods used are indicated in the appropriate figure legends. In brief, 1-way ANOVA followed by Tukey’s multiple-comparison test was used for the statistical analysis of the results presented in Fig 4 and 5. 2-way ANOVA followed by Tukey’s multiple-comparison test was used for (Fig 1, 3, 6, SI4 and SI6). T-test was used for the statistical analysis of the results presented in Fig 2, 3, 5, 7 and SI5. Mann-Whitney u-test was used for the statistical analysis of the results presented in Fig 5 and SI5. The significance level was set at P less than 0.05: *P < 0.05; **P < 0.01; ***P < 0.001; ****P < 0.0001.

**Fig 1:**
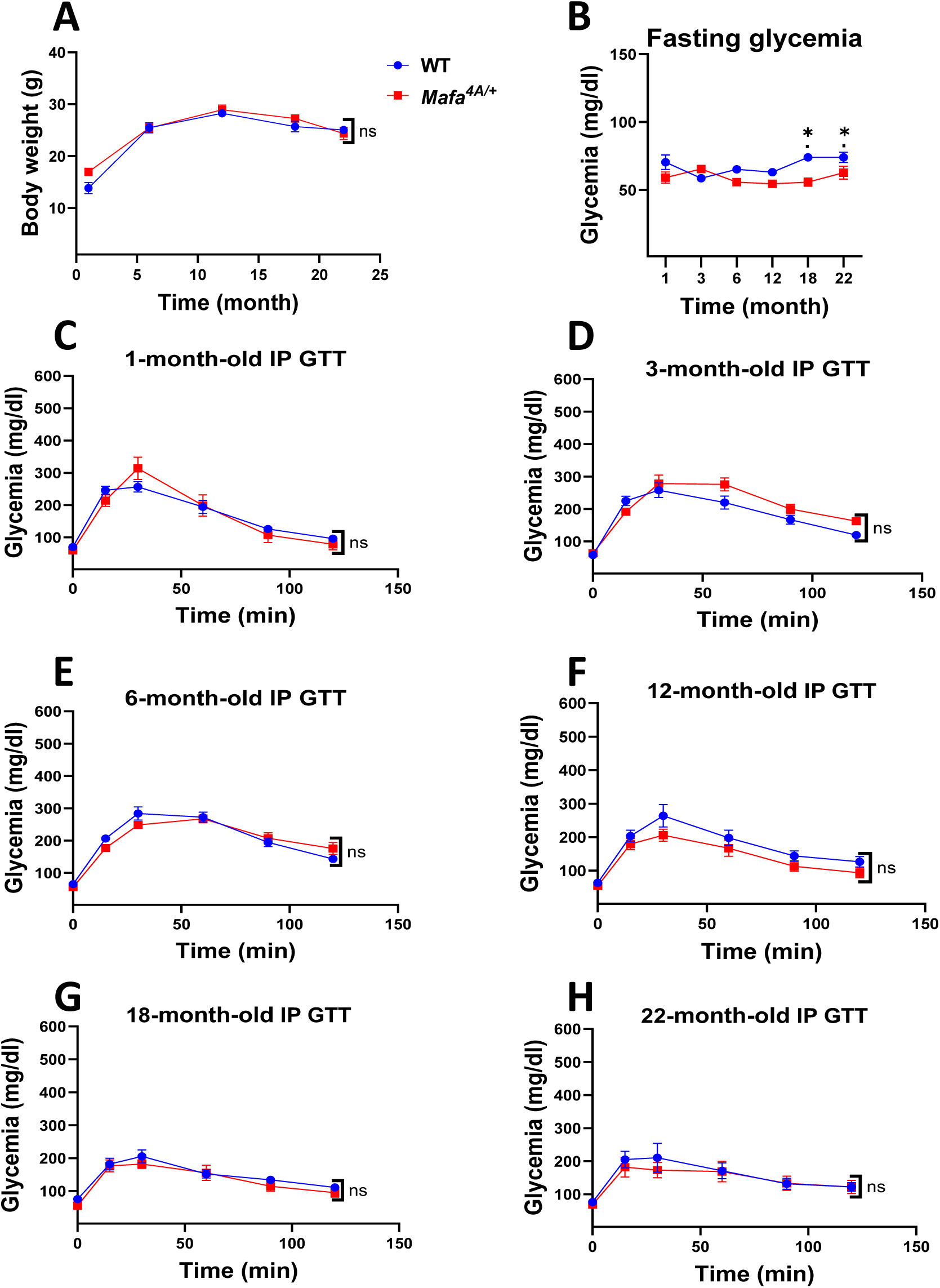
Physiological and metabolic analysis of WT and *Mafa^4A^*^/+^ male mice. A) analysis of weight gain from 1 to 22 months (*n>*3). B) Fasting blood glucose of mice from 1 to 22-month-old after fasting for 16 hours (*n>5*). C-H) Intraperitoneal glucose tolerance test (IP GTT) performed after 16 hours of fasting on mice from 1 to 22 months (*n>7*). Data represent a mean ± SEM of the *n. 2*-way ANOVA followed by Tukey’s multiple-comparison test was used. *p<0.05.

**Fig 2:**
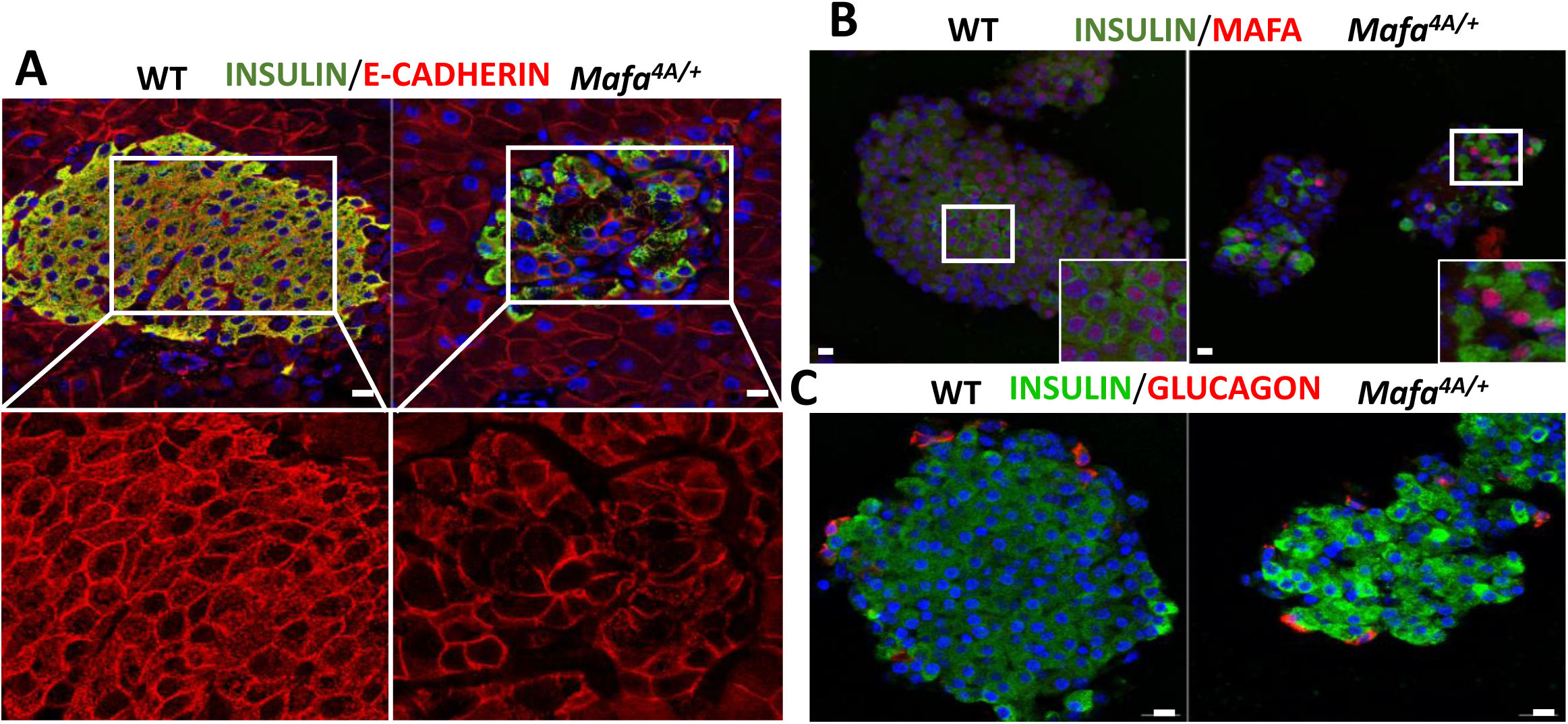
Histological analysis of the pancreases from 4-month-old *Mafa^4A^*^/+^ and WT male mice. A) Immunostaining with anti-insulin (green) and anti-E-cadherin (red) antibodies. Higher magnification of the E-cadherin staining in the bottom. B) Isolated pancreatic islets were stained for MAFA in red and insulin in green. C) Isolated pancreatic islets were stained for insulin in green and for glucagon in red to detect beta and alpha cells respectively. Scale at 10μm

**Fig 3:**
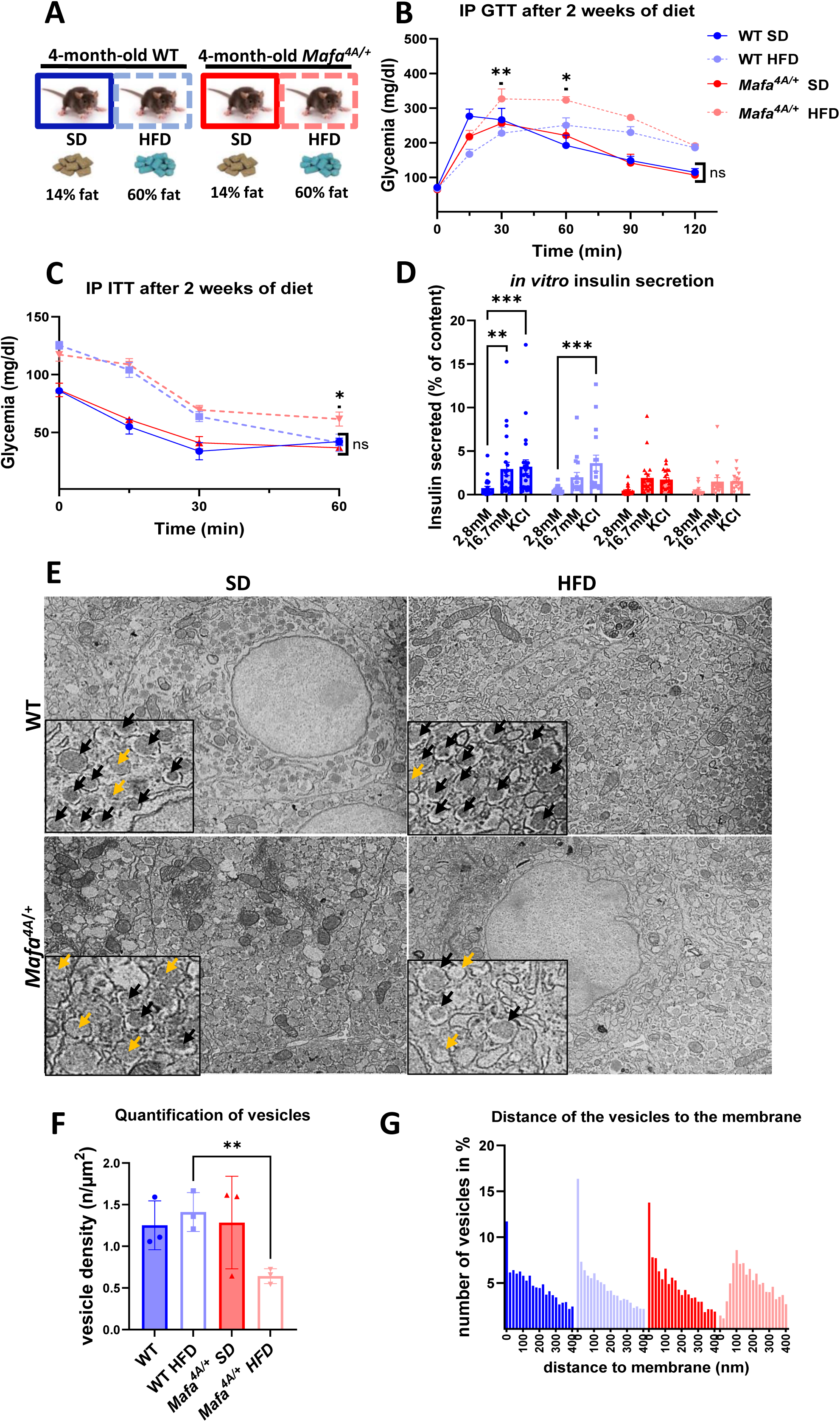
Analysis of the effects of high fat diet on male mice. A) experimental design. SD: standard diet, HFD: High Fat Diet. B) Intraperitoneal glucose tolerance test (IP GTT) (*n>*3). C) Intraperitoneal insulin tolerance test (IP ITT) (*n>*3). D) Glucose stimulated insulin secretion (GSIS) on islets isolated from WT and mutant mice. Secreted insulin was measured by ELISA and related to the total content of insulin (*n*>16). E) Transmission electron microscopy images showing the ultrastructure of pancreatic beta cells, highlighting insulin secretory granules: mature granules (black arrow) with a dense core and clear halo, and immature granules (yellow arrow) with lower electron density and lacking a halo (magnification ×2500). F-G) Quantification of the number of secretory granules (F) and their distance to the membrane (G) by deep learning (*n=*3). Data represent a mean ± SEM of *n*. *2*-way ANOVA followed by Tukey’s multiple-comparison test was used for B-C-D and unpaired t-test for F. *p<0.05; **p<0.01; ***p<0.001.

**Fig 4:**
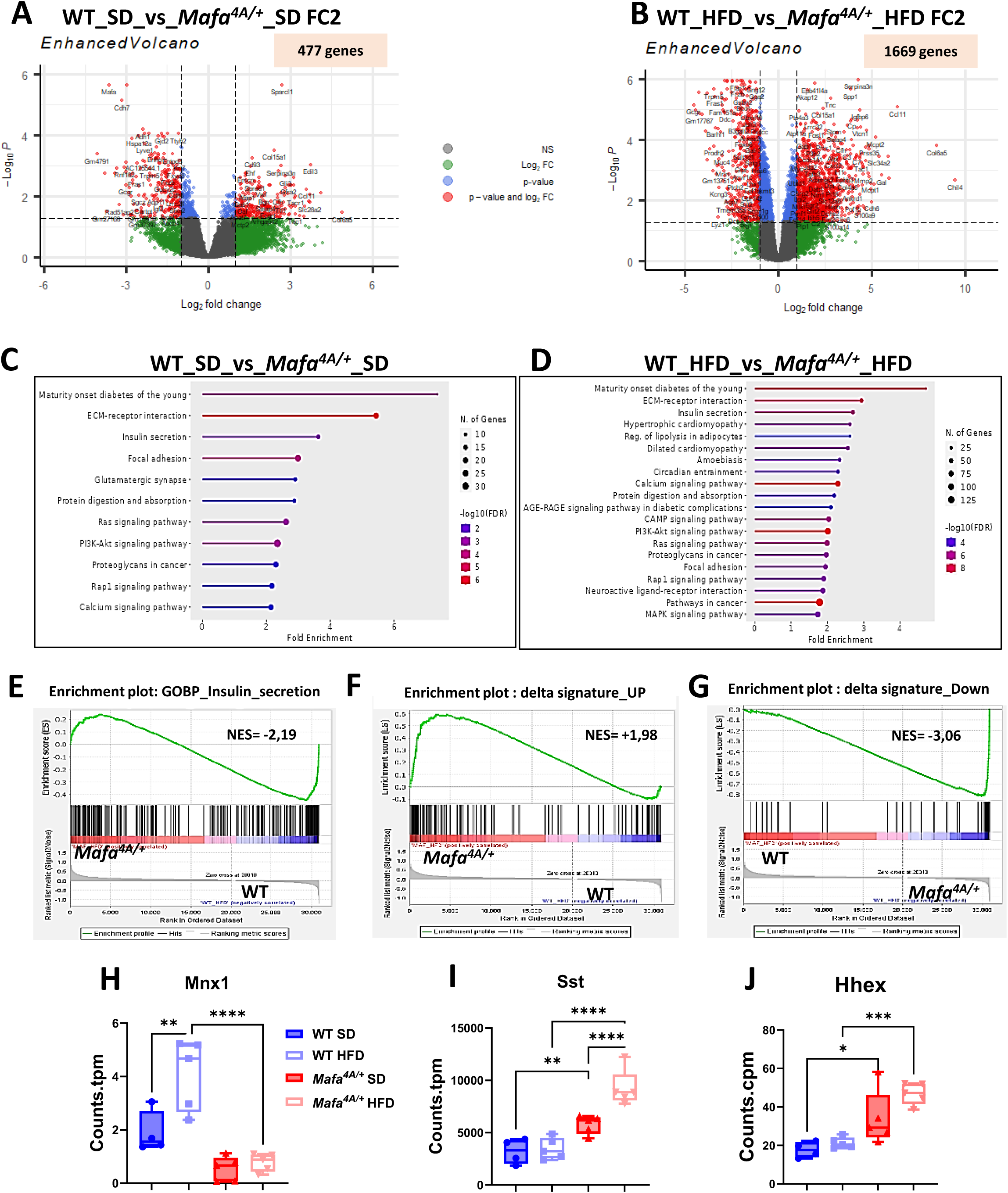
Gene expression analysis by bulk RNAseq. A-B) Volcanoplot. C-D) Diagram showing a strong signature of certain pathways (MODY and insulin secretion) in mutants as compared to WT controls under SD or HFD. E-G) Enrichment curve obtained by GSEA showing the variations in the expression of genes involved in insulin secretion (E) and in delta cell identity (F and G). H-J) Box-plot representation of the expression level of the *Mnx1*, Somatostatin (*Sst*) and *Hhex* genes in mice. Data represent a mean ± SEM of *n. 1*-way ANOVA followed by Tukey’s multiple-comparison test was used. *p<0.05; **p<0.01; ***p<0.001; ****p<0.0001.

**Fig 5:**
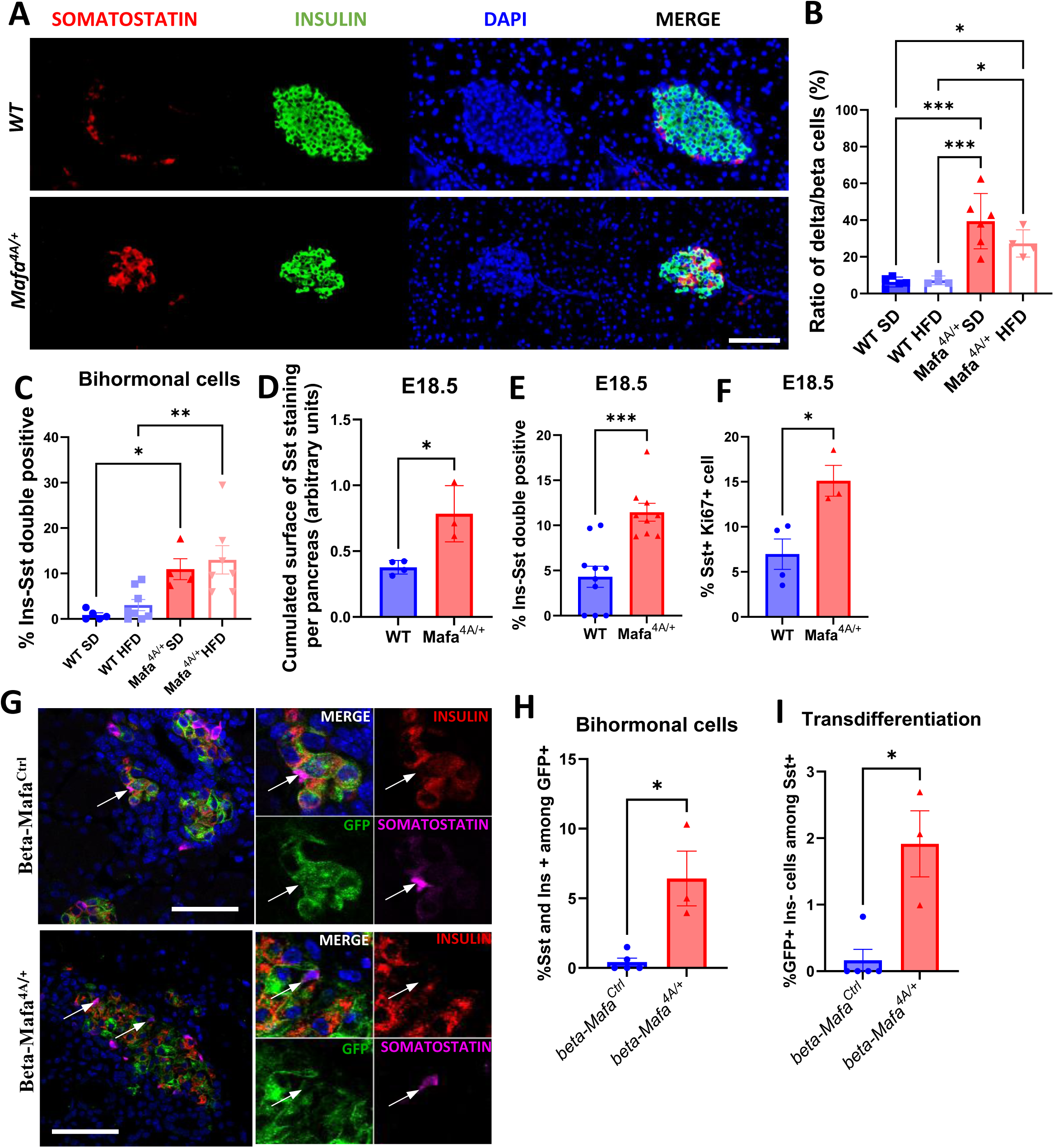
Increased number of delta cells in mutant mice. A) Immunofluorescent detection of beta cells with an anti-insulin antibody (green) and delta cells with an anti-somatostatin antibody (red). B-C) Quantification of delta cells (B) and insulin-somatostatin bihormonal cells (C) in the islet of WT and mutant mice exposed to SD or HFD (*n>4*). D-F) Quantification of the absolute numbers *Sst* expressing cells at E18.5 (D), quantification of the percentage of insulin-somatostatin bihormonal cells among insulin expressing cells (E) and quantification of the proliferation of delta cells (percentage of Ki67+/insulin+ cells) (F) (*n>*3). G) Immunostaining of beta cells with an anti-insulin antibody (red), delta cells with an anti-somatostatin antibody (pink) and GFP with anti-GFP antibody (green) at E18.5 in pancreas of beta-*Mafa^4A/+^* and beta-*Mafa^Ctrl^*. Arrows indicate cells that are positive for both Somatostatin and GFP. H-I) Quantification of insulin-somatostatin bihormonal cells (H) and somatostatin-GFP positive but insulin negative (I) (*n>*3). This combination represents beta to delta transdifferentiated cells. Data represent a mean ± SEM of *n*. 1-way ANOVA followed by Tukey’s multiple-comparison test was used in B-C, unpaired t-test for D-E-F and Mann-Whitney test for G-H. *p<0.05; **p<0.01; ***p<0.001; ****p<0.0001.(60μm scale)

### RNAseq

Total RNA was prepared from pancreatic islets isolated as described above using the Rneasy mini kit (QIAGEN). Either 10 or 4 ng of total RNA was used for the construction of sequencing libraries. RNA libraries were prepared using the Takara SMARTer Stranded Total RNA-Seq Kit-Pico Input Mammalian kit following the manufacturer’s protocols. Sequencing was carried out on a NovaSeq 6000 instrument from Illumina (S1-PE100, paired-end reads). Raw sequencing reads were first checked for quality with Fastqc (0.11.9) and trimmed for adapter sequences with cutadapt (1.12) using the TrimGalore (0.6.7) wrapper. Trimmed reads were subsequently aligned to the complete human ribosomal RNA sequence with bowtie (1.3.0). Reads that did not align to rRNA were then mapped to the mouse reference genome mm10 and read counts per gene were generated with STAR mapper (2.7.6a). The bioinformatics pipelines used for these tasks are available online (RNAseq v4.0.4: https://gitlab.curie.fr/data-analysis/RNA-seq, doi: 10.5281/zenodo.7443721). Counts were normalized using TMM normalization from EdgeR (v 3.32.1). Differential gene expression was assessed with the Limma voom framework (v 3.46.0). Genes with an absolute fold-change ≥1.5 and an adjusted p-value<0.05 were labeled significant.

### Functional Analyses

Gene Ontology analyses were performed by using R (4.0.3). Gene Sets Enrichment Analyses (GSEA, http://software.broadinstitute.org/gsea) were run using signal-to-noise for the ranking gene metric and 1000 permutations. The DELTA signature used as gene set reference was obtained from Gribble et al, 2007 (27) by sorting genes based on FC with FC < -2 for down genes and FC > 2 for up genes in both delta_vs_beta and delta_vs_alpha conditions.

## RESULTS

### *Mafa^4A^* mutation leads to lifelong MAFA protein stabilization

In our previous work (4), we showed that preventing GSK3-mediated MAFA phosphorylation leads to its stabilization. Thus, we first checked if MAFA is stabilized in the pancreas of *Mafa^4A/+^* mice. We performed IF analysis using anti-insulin and anti-MAFA antibodies. We indeed showed that the MAFA signal was strongly increased in beta cells of knock-in mice at the fetal-(E18.5), the adult-stage (3, 6 months) as well as old mice (22 months) (Fig. SI1-3) both in male and female. Next, we investigated the phenotype of these mice in a sex-dependent manner.

### Glucose homeostasis in *Mafa^4A/+^* males

We first analyzed the phenotype of *Mafa^4A^* ^/+^ males. The body weight (Fig. 1A), fasting blood glucose (Fig. 1B), glucose and insulin tolerance tests (GTT, Fig. 1C-H and ITT, Fig. SI4) of *Mafa^4A^*^/+^ males were mostly normal as compared to WT animals, although a mild hypoglycemia was observed in old animals (Fig. 1B). In striking contrast, the islet architecture was significantly disorganized in adult mutant mice (Fig. 2 A-C). In particular, the presence of E-cadherin at the membrane was decreased, and beta cell-cell contacts were strongly altered (Fig. 2A). The localization of glucagon-producing alpha cells at the islet periphery was normal. The pancreas weight of the mutants and the controls was similar (203 +/- 20 mg versus 183 +/- 7 mg respectively, ns, n>6) and the quantification of alpha and beta cell masses did not differ from that of controls (Fig. SI5). We next explored the sensitivity of *Mafa^4A^*^/+^ male mice to the nutritional environment. Mice were fed with a high-fat diet (HFD) for 2 weeks. Male *Mafa^4A^*^/+^ mice treated with HFD developed significantly more pronounced glucose intolerance than WT-HFD mice (Fig.3. A-B). While ITT was altered in HFD-fed mice, no difference in insulin sensitivity in *Mafa^4A^*^/+^ HFD mice compared to WT-HFD control mice was observed (Fig. 3C). Together, these data suggest that *Mafa^4A^*^/+^ HFD mice are prone to beta-cell dysfunction. To test this hypothesis, we analyzed GSIS from pancreatic islets isolated from *Mafa^4A^*^/+^ SD (standard diet), *Mafa^4A^*^/+^ HFD, WT-SD, and WT-HFD mice (Fig. 3D). In control islets (WT-SD), insulin secretion was stimulated by both high glucose and KCl as expected. HFD exposure decreased the response to high glucose but not to KCl. In contrast, GSIS was dramatically impaired in *Mafa^4A^*^/+^ mice. Together, our *in vivo* and *ex vivo* data show that the *Mafa^4A^* mutation leads to beta-cell defects that worsens with HFD.

### Pancreatic beta cell impairment in *Mafa^4A^*^/+^ male mice

To decipher the origin of GSIS alteration in our mutant mice, we analyzed the ultrastructure of beta cells using an electron microscope (Fig. 3E). In WT-SD and WT-HFD mice, the characteristics of secretory granules were normal (black arrows). However, in *Mafa^4A^*^/+^ SD mice, secretory granules displayed an immature appearance (yellow arrows) with a lack of dense nuclei or empty contents. In *Mafa^4A^*^/+^ HFD mice, this phenotype was more pronounced. Using deep learning, we quantified vesicle density and their distance from the membrane. The latter is indicative of insulin secretion capacity. We found that in *MafA^4A/+^* HFD islets, the total number of granules and those that are close to the membrane (<100 um) were significantly reduced compared to the other groups (Fig. 3F-G). Thus, these results indicate that the secretion mechanism is disrupted in *Mafa^4A^*^/+^ mice fed with HFD.

To better understand the mechanisms of this dysfunction, we performed bulk RNA sequencing analysis on islets extracted from the 4 different groups of mice, indicating a strong differential expression of genes (Fig. 4A-B). Pathway analyses was performed on DEG (differentially expressed genes) in islets from *Mafa^4A^*^/+^ vs WT mice in different diet conditions. We found a Maturity Onset Diabetes of the Young (MODY) signature enriched in *Mafa^4A^*^/+^ SD and HFD islets as compared to corresponding controls, as well as impaired insulin secretion and calcium signaling (Fig. 4C-D). Specifically, DEG analysis showed that *Mafa^4A^*^/+^ SD and HFD islets have decreased expression of (i) a number of important genes that determine beta cell identity (*Iapp*, *Ins1*, *Ins2*, *Isl1*, *Mafa*, *Mnx1*, *Nkx2-2*, *Pax6*, *Pdx1*, *Tshz1*), (ii) genes associated with metabolism (*Eno2*, *Glp1r*, *Igf1r*, *Slc2a2* [Glut2], *Slc30a8* [Znt8], *Ucn3*), (iii) genes involved in secretion (*Atp2a2*, *Atp2b2*, *Bace2*, *Ncald*, *Rims3*, *Syt13*, *Syt7*, *Wfs1*) (Fig. 4E), and (iv) increased expression of prohibited genes (*Fcgrt*, *LdhA*, *Oat*, *Yap1*, *Zcchc24*). The concept of forbidden genes was first described by Rutter et al., showing that shutting down a subset of disallowed genes is important for the differentiated function of beta-cells, and for the control of beta-cell expansion and insulin secretion (28). For most of these genes, the variation in their expression is exacerbated by HFD. Importantly, we also observed a marked decrease in *Mnx1* expression in *Mafa^4A^*^/+^ mice (Fig. 4H), a reduction previously linked to beta-cell to delta-cell transdifferentiation (29). We thus investigated enrichment of the delta-cell signature both in the SD and HFD conditions (described in (27)) (Fig. 4F-G and I-J).

### The development of delta cells is amplified in the *Mafa^4A/+^* male mice

We found an enriched signature of the delta-cell program in the islets from the adult *Mafa^4A/+^* SD mice (Fig. 4F-G) and in particular an increased expression of the somatostatin (*sst*) and *Hhex* genes (Fig. 4I-J). This increased expression was even more significant in the HFD condition. These results suggest an increase of delta-cells when *Mafa^4A^* is present. To further characterize this phenotype, we performed IF experiments using anti-insulin and anti-somatostatin antibodies (Fig. 5A). We observed a mislocalization of somatostatin-positive cells in *Mafa^4A^*^/+^ animals (Fig. 5A) and an 8-fold increase (39% in the mutants versus 5% in the controls, p<0.001) of the percentage of delta-cells in the mutants (Fig. 5B). A similar difference between the mutant and control mice was observed with HFD. We also detected some bi-hormonal cells that co-expressed insulin and somatostatin (Fig. 5C). Indeed, in WT SD and WT HFD islets, such cells were scarce, reaching respectively 1.2% and 2.1% of the insulin-positive cells. On the contrary, in the *Mafa^4A^*^/+^ SD and *Mafa^4A^*^/+^ HFD, the percentage of insulin-positive cells that co-expressed somatostatin was 10.7% and 13% respectively. Altogether, these results suggest a trans-differentiation of beta-cells to delta-cells. To determine whether this event has a developmental origin, we analyzed the development of beta- and delta-cells at the fetal stage E18.5. Indeed, at this stage, *Mafa* is already expressed (Fig. SI1) and the segregation between beta- and delta-cells is expected to have already occurred (21).

At E18.5, the size of WT and *Mafa^4A/+^* male pancreases were not statistically different (cumulated surface of 1500 +/-106 AU in WT versus 2196 +/- 905 in the mutants, n>3, p=0.41). In the WT controls we were only able to find rare-delta-cells and a few insulin-expressing cells co-expressing somatostatin (4% of the insulin-expressing cells) (Fig. 5D-E). In contrast, the number of delta cells increased in the mutants, due at least partly to the upregulation of delta-cell proliferation (Fig. 5F). Moreover, the number of insulin^+^/somatostatin^+^ cells was significantly increased in the mutants as compared to the controls (11% of insulin^+^/somatostatin^+^ among insulin-expressing cells in the mutants versus 4% in the controls) (Fig. 5E). Thus, these data indicate that the bi-hormonal beta/delta cells arise during embryogenesis. To further research whether the beta/delta cells result from the trans-differentiation of *Mafa^4A^*-expressing-beta cells, we performed a lineage-tracing analysis (Fig. 5G). For this, we generated pIns1-Cre^ERT2^, mT/mG, *Mafa^4Afl/+^* mice (beta-*Mafa^4A^*^/+^). We also used pIns1-Cre^ERT2^, mT/mG, *Mafa^+/+^* as controls (beta-*Mafa^Ctrl^*). We treated pregnant mice of each group with tamoxifen at E13.5 to activate the recombinase and we next analyzed the pancreases of the fetuses at E18.5. In both beta-*Mafa^4A^*^/+^ mutant and the beta-*Mafa^Ctrl^* mice (Fig. 5G), most of the insulin expressing cells were also positive for GFP as expected, showing the efficient recombination of the tomato (mT/mG) locus in beta-cells. Thus, the GFP reporter gene can be used for the tracing of beta-cells and their descendants. In mice carrying the *Mafa^4A^* mutation, we found a significant increase of the number of cells that co-express GFP, insulin and somatostatin as compared to the control mice (Fig. 5H). This result indicates that beta-cells that express insulin and *Mafa^4A^*, also express somatostatin. We also quantified the percentage of the somatostatin-positive cells that are positive for GFP but negative for insulin. These cells initially exhibit a beta-cell identity (GFP⁺) but later adopt a delta-cell phenotype (Ins⁻/Sst⁺), likely reflecting a transdifferentiation process. We found an 18-fold increase of these cells in the beta-*Mafa^4A^*^/+^ mice vs the controls (Fig. 5I). Collectively, these findings provide strong evidence for beta-to-delta cell transdifferentiation upon Mafa^4A^ expression in beta-cells. This conversion appears to occur during fetal development and may alternatively reflect aberrant differentiation of beta- and delta-cells originating from a shared progenitor. To further investigate the sex-dependent roles of *Mafa^4A^*, we next conducted our study on female mice.

### *Mafa^4A^*^/+^ female mice are hypoglycemic

We examined *Mafa^4A^*^/+^ female mice from 1 to 22 months of age. Their growth did not differ from control mice (Fig. 6A). Surprisingly, their fasting glucose was decreased compared to control mice and this difference, being significant at 3, 6, 18 and 22 months, was more pronounced during aging (Fig. 6B). This hypoglycemia was observed after 6 months when GTT experiments were performed (Fig. 6 C-H). To better understand this phenotype, we first performed ITT analysis (Fig. SI6A-F), and we measured serum insulin levels in *Mafa^4A^*^/+^ female mice at 22 months of age under fasting and refeeding conditions (Fig. SI6G). We found no difference between mutant female mice and controls. The GSIS of islets isolated from *Mafa^4A^*^/+^ female mice also did not differ from that of controls (Fig. SI6H-I). We therefore concluded that the hypoglycemia that we observed was unlikely to be caused by beta-cell overactivity.

**Fig 6:**
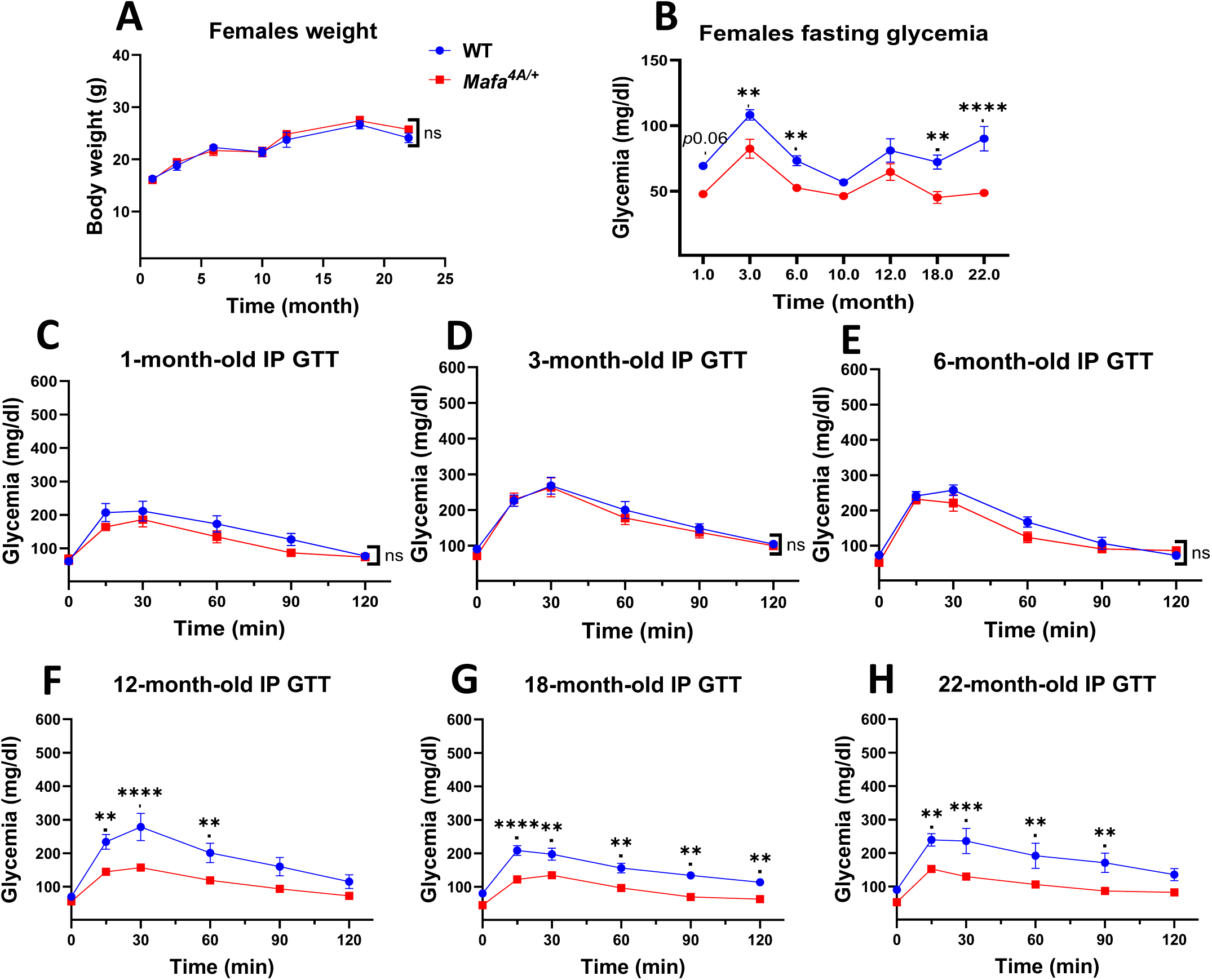
Physiological and metabolic analysis of *Mafa^4A/+^* female mice. A) Analysis of weight gain from 1 to 22 months (*n>5*). B) Fasting blood glucose of mice from 1 to 22 months (*n>5*). C-H) Intraperitoneal glucose tolerance test (IP GTT) (*n>6*). Data represent a mean ± SEM of *n*. 2-way ANOVA followed by Tukey’s multiple-comparison test was used. *=p<0.05; **p<0.01; ***p<0.001; ****p<0.0001.

### Decreased beta-cell mass in *Mafa^4A^*^/+^ female mice

The total pancreatic weight of the mutants was similar to that of the controls (212 +/- 33 mg in the WT pancreases versus 166 +/- 8 mg in the mutants, n=3, p=0,25). Since the islet architecture was altered in *Mafa^4A^*^/+^ male mice, we investigated female islet histology by IF in females with anti-insulin and anti-glucagon antibodies to detect beta- and alpha-cells, respectively (Fig. 7A). A significant reduction in beta-cell mass was found in 3-month-old female islets (Fig. 7B). Additionally, the number of alpha-cells was slightly but not significantly increased in *Mafa^4A^*^/+^ female mice (Fig. 7C).

**Fig 7:**
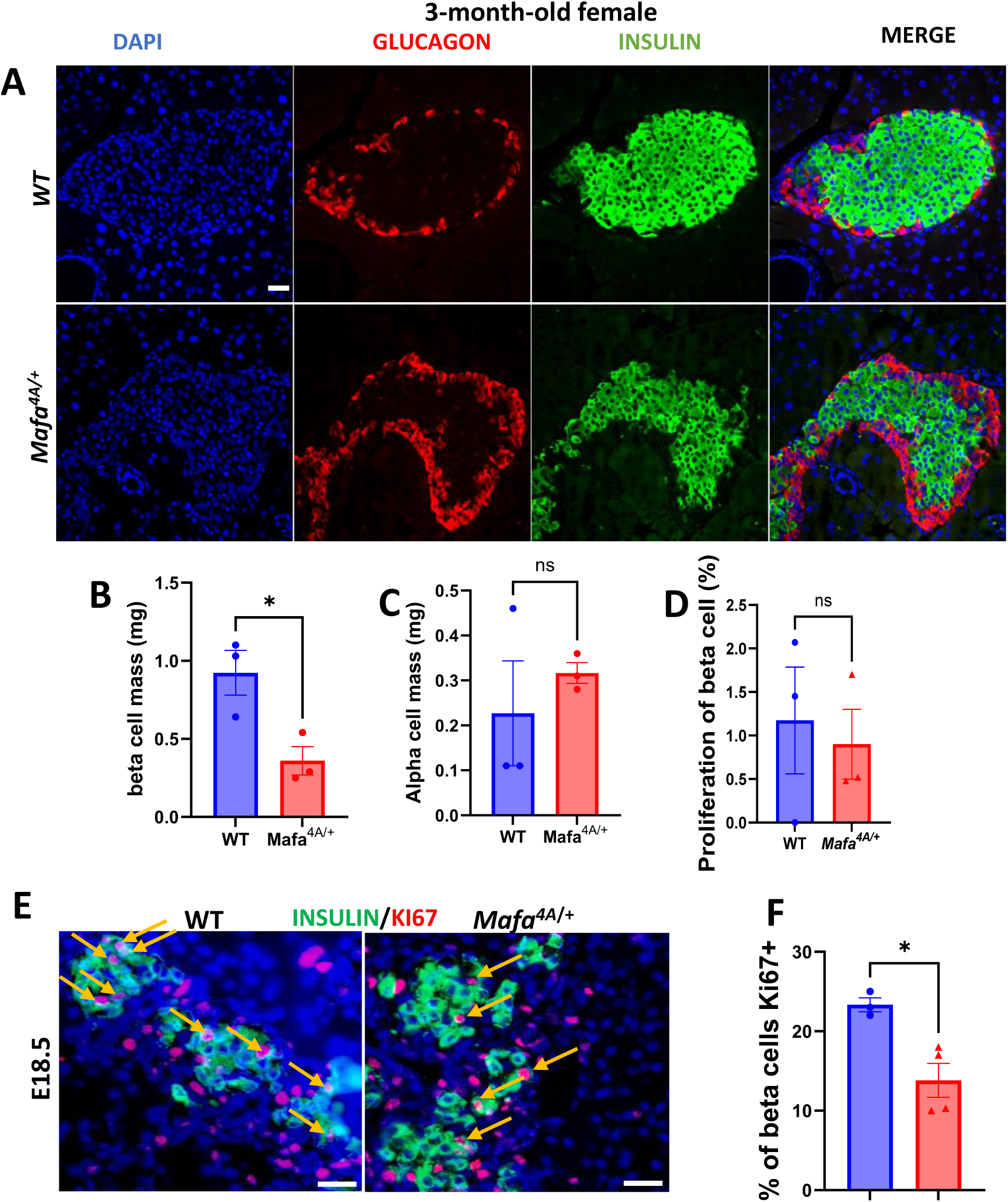
Histological analysis of the pancreases of female mice. A) Beta and alpha-cells were detected respectively with an anti-insulin antibody (green) and with an anti-glucagon antibody (red). B-D) Quantification of beta cell mass (B), alpha cell mass (C) and proliferation of beta cells (D) (*n=*3). E) Evaluation of the beta cell proliferation at E18.5 by immunofluorescence using an anti-insulin antibody in green and an anti-Ki67 antibody in red. F) Quantification of the beta cell proliferation (*n>*3). Data represent a mean ± SEM of *n*. Unpaired t-test was used for B-C-D-F. *p<0.05. Scale at 20μm

To investigate the cause of the reduced beta-cell mass, we analyzed beta-cell proliferation by IF at 3 months, using anti-insulin and anti-Ki67 antibodies. No difference was found (Fig. 7D). We then examined whether a developmental origin could be responsible for the decrease of beta-cell mass. We dissected pancreases at E18.5, an important time point of beta-cell proliferation. We found a two-fold reduction in beta-cell proliferation in *Mafa^4A^*^/+^ female fetuses compared to controls (Fig. 7E-F). Thus, these data show that GSK3-mediated MAFA phosphorylation plays an important role in fetal beta-cell proliferation in females, thereby controlling the beta-cell pool.

### Enlarged cystic inflammatory pancreatic ducts in aged *Mafa^4A^*^/+^ female mice

In the pancreases of 22-24 months old *Mafa^4A^*^/+^ females, we found 8 ductal inflammatory dilatations (8/43) and 3 others corresponding to typical mucinous cystic neoplasia (MCN) (3/43) (Fig. 8A). Indeed, they were characterized by the presence of pseudo-ovarian stroma and mucinous cells (Fig. 8B). These cysts stained positive with alcian blue, revealing the presence of mucin. The borders of the cysts were positive for SOX9 (Fig. 8C), and inflammation was revealed by the presence of immune cells as well as F4/80 staining (Fig. 8D). On the contrary, these structures were never found in the age-matched WT female mice nor the males (WT and *Mafa^4A^*^/+^). Altogether, these data indicate that *Mafa^4A^*expression renders the females prone to develop cystic inflammation and MCNs, that represent a precursor step to pancreatic ductal adenocarcinoma (PDAC). To investigate the clinical relevance of these results, we probed for the presence of MAFA in MCNs biopsies from patients histologically and we observed the presence of MAFA-expressing cells (Fig. 8E), suggesting that MAFA could be a potential novel marker of human MCNs.

**Fig 8:**
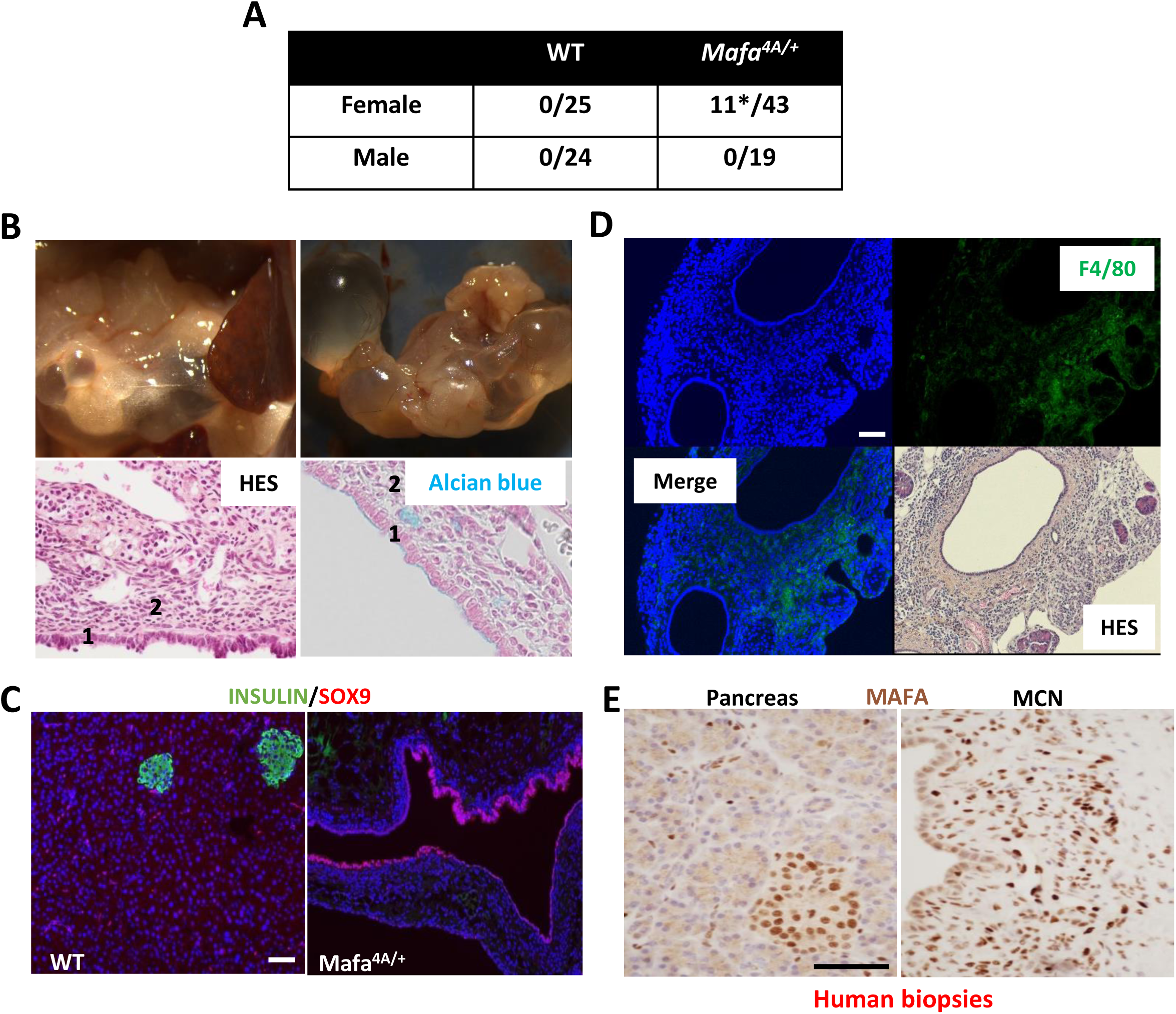
Histological analysis of female mice. A) Quantification of the cystic dilatations observed in mice at 22 months. Of note 3 among 11 were MCNs. B) Image of mouse pancreas at 22 months showing the presence of cysts before sampling (top left) and after sampling (top right). A hematoxylin eosin staining shows the presence of a unistratified epithelium (1) and a pseudo-ovarian stroma (2) (bottom left) and a staining with alcian blue shows the presence of mucin on the epithelium (bottom right). C) Immunostaining of beta cells with an anti-insulin antibody in green and SOX9, a characteristic marker of ductal cells, in red. D) Detection of inflammation by immunostaining of macrophages with an anti-F4/80 antibody and by HES. Characteristic micronucleated cells can be visualized. E) Immunostaining of MAFA in the stroma of human neoplastic mucinous cysts (MCNs) of the pancreas, and in a human pancreas sample. Scale at 60μm

## DISCUSSION

In the present study, we show that GSK3-mediated phosphorylation of MAFA is a key determinant of beta-cell identity in the male mice. *Mafa^4A^*^/+^ mice are more prone to diabetes when the nutritional environment is modified to contain high dietary fat. In particular, the machinery of insulin secretion is altered when such phosphorylations are prevented.

Using Transmission electron microscopy (TME), we showed that the defect of GSIS is associated with a decreased number of insulin granules and their immaturity. This observation is also supported by our RNAseq data showing the deregulation of many genes involved in secretion. On the other hand, the islet architecture was also impaired. Recently, Cottet-Dumoulin et al. showed that beta-cell-cell contacts favor insulin secretion and protein biosynthesis at low or moderate glucose concentrations (30). At high glucose concentrations, insulin secretion is also increased by these contacts but not protein biosynthesis. The abnormal contacts between the *Mafa^4A^*^/+^ beta cells (Fig. 2A) may thus contribute to the alteration of insulin secretion.

Importantly, we also identified a strong modification of beta-cell identity in *Mafa^4A^*^/+^ males with beta- to delta-cell transition observed. In previous studies (31), the implication of *Mafa* in beta-cell plasticity was also highlighted. Indeed, a beta-to-alpha cell trans-differentiation was found in the *Mafa* knock-out mice while a beta-to-delta cell trans-differentiation is observed in our *Mafa^4A^*^/+^ knock-in mice. This indicates that not only the presence of *Mafa* is important, but also its phosphorylation by GSK3 is essential to maintain a correct beta-cell identity. This additionally suggests that physiological MAFA activity prevents endocrine plasticity by strengthening beta-cells identity. It is currently unclear why a complete loss of *Mafa* induces a shift from beta-to alpha-cells, while preventing its phosphorylation induces a shift from beta-to delta-cells. Moreover, hormonal regulations may also contribute to abnormal GSIS. Indeed, somatostatin is a well-established inhibitor of insulin secretion. The increased development of delta-cells in our *Mafa^4A^*^/+^ mice is likely to increase somatostatin signaling within islets, and in consequence, to negatively regulate GSIS. Therefore, defects of insulin secretion in male *Mafa^4A^*^/+^ animals appear to be multifactorial, involving defects in the secretory machinery directly by deregulation of key genes, but also indirectly due to alteration of beta-cell-cell contacts as well as hormonal impairment.

Our work also provides insight into the extent to which impaired MAFA phosphorylation contributes to the S64P syndrome observed in multiple families with diabetes and insulinoma (17). Walker et al. introduced the S64F mutation (18) in mice. This mutation is close to the S65 phosphorylation, a priming site that controls GSK3-mediated phosphorylation and ubiquitin-mediated degradation of MAFA. These S64F mice showed glucose intolerance in males that was preceded by transiently higher MAFA protein levels at 4 weeks, but not later. These results suggest that at least some of the effects of the S64F mutation involve MAFA stabilization. In contrast, we found that the *Mafa^4A^* mutation leads to a permanent stabilization of MAFA protein in the beta-cells both in males and females. We therefore believe that the molecular mechanisms observed in the *Mafa^4A^*^/+^ mice, although mostly similar to those described in the S64F mice, may differ in certain aspects. Indeed, the effects of the *Mafa^4A^* mutation are probably more direct on the phosphorylation and stabilization of MAFA, and it cannot be excluded that the S64F mutation involves additional mechanisms on the transcriptional activity of MAFA that were not yet explored.

In female mice, we did not investigate the effects of HFD. Indeed, it is well-established that female mice are protected against HFD-induced metabolic syndrome (32). Accordingly, GSIS by the female islets were normal (Fig SI6), showing that these islets were not affected in the same way as compared to the males. Surprisingly, unlike males, the *Mafa^4A^*^/+^ females were hypoglycemic. This result was not expected, and this effect of MAFA should be further investigated in future analysis.

Interestingly, we found a decreased proliferation of the beta-cells in the *Mafa^4A^*^/+^ female fetal mice. Previously, other studies also showed that *Mafa* deletion leads to decreased expression of the prolactin receptor and its target, cyclin D2 (33). Consistently, the loss of *Mafa* resulted in decreased beta-cell proliferation at 4 weeks. Our observations with *Mafa^4A^*were similar. However, the timing of the reduction of proliferation in the *Mafa^4A^*^/+^ mice differs from the *Mafa* knock-out mice. In addition, our results displayed a sex bias not described in KO mice. Some of these differences likely stem from the distinct nature of these mutations (Knock-Out versus *Mafa^4A^*), but the sex bias in the *Mafa^4A^*^/+^ mice remain unexplained. Interestingly, a sex-biased phenotype was also observed in the *S64F* mice (18), again highlighting some similarities between the two models. The specific phenotype of females was not affected by ovariectomy, suggesting that estrogen sex hormones were not involved.

Interestingly, we found pancreatic cystic structures in the *Mafa^4A^*^/+^ aging females. Some of them were identified as mucinous cystic neoplasms (MCNs). To date, the causes of these lesions in humans remain largely unknown. The presence of the MAFA protein in biopsies of MCN patients suggests that MAFA may have a role in the development of these tumors. Interestingly in humans, 95% of MCNs are found in females. Similarly, MCNs were detected only in our aging female mice but not in males, suggesting that the *Mafa^4A^*^/+^ mice represent a good model for this disease. Recent data also showed that aging is a key determinant of both endocrine and exocrine tumors (34). In particular, the age-related morphological changes in the pancreas are associated with carcinogenesis (35) and gradual telomere shortening also occurs in the tumorigenesis of MCNs (36). Moreover in our samples, we also detected signs of inflammation and lipogenesis, that probably participate to the formation of neoplasms (37).

In conclusion, our data demonstrates the crucial role of GSK3-mediated phosphorylation of MAFA in the determination of the functional identity of pancreatic beta-cells. It also suggests the implication of MAFA in cystic neoplasms that represent precursors of PDAC. To date, no therapeutic strategies have been developed to directly target MAFA, but we remain hopeful that advances in pharmacological research will allow for the discovery of new treatments for patients suffering from diabetes (MODY) or pancreatic neoplasms.

## Acknowledgements

We thank NG Dave YX for proofreading our manuscript.

