## Supplemental Tables and Figures for "MAFA Phosphorylation Controls Beta-Cell Identity and Sex-Specific Pancreatic Disease Outcomes"

| TABLE 1 primary antibody | | | | | | |
|---|---|---|---|---|---|---|
| Protein | ref Provider | Provider | species | against | experiments | Dilution |
| INSULIN | 20056 | immunostar | RABBIT | MOUSE | IF | 1/1000 |
| INSULIN | I2018 | SIGMA | MOUSE | MOUSE | IF | 1/1000 |
| GLUCAGON | 6728-1-IG | PROTEINTECH | MOUSE | MOUSE | IF | 1/1000 |
| MAFA | D2Z6N | CELL SIGNALING | RABBIT | MOUSE | IF/WB | 1/100 |
| Ki67 | 550609 | BD BIO | MOUSE | MOUSE | IF | 1/200 |
| SOMATOSTATIN | SC-74556 | SANTA CRUZ | MOUSE | MOUSE | IF | 1/1000 |
| SOX9 | AB5535 | MILLIPORE MERCK | RABBIT | MOUSE | IF | 1/1000 |
| F4/80 | AB6640 | ABCAM | RAT | MOUSE | IF | 1/500 |
| E-CADHERIN | 610182 | BD BIO | MOUSE | MOUSE | IF | 1/100 |
| GFP | AB13970 | ABCAM | CHK | MOUSE | IF | 1/1000 |

| TABLE 2 secondary antibody | | | | | |
|---|---|---|---|---|---|
| species | Protein | ref Provider | Provider | species | against |
| Chicken | Alexa goat anti chicken 488 | A11039 | Molecular probes/invitrogen | goat | chicken |
| Rabbit | Alexa goat anti rabbit 488 | A11008 | Molecular probes/invitrogen | goat | rabbit |
| | Alexa goat anti rabbit 555 | A21428 | Molecular probes/invitrogen | goat | rabbit |
| | Alexa goat anti rabbit 594 | A11037 | Molecular probes/invitrogen | goat | rabbit |
| Rat | Alexa chicken anti rat 488 | A21470 | Molecular probes/invitrogen | chicken | rat |
| Mouse | Alexa goat anti mouse 647 | A32728 | Molecular probes/invitrogen | goat | mousse |
| | Alexa goat anti mouse 488 | A21042 | Molecular probes/invitrogen | goat | mouse |
| | Alexa goat anti mouse 555 | A21422 | Molecular probes/invitrogen | goat | mouse |
| | Alexa donkey anti mouse 555 | A21203 | Molecular probes/invitrogen | donkey | mouse |

### Fig SI 1

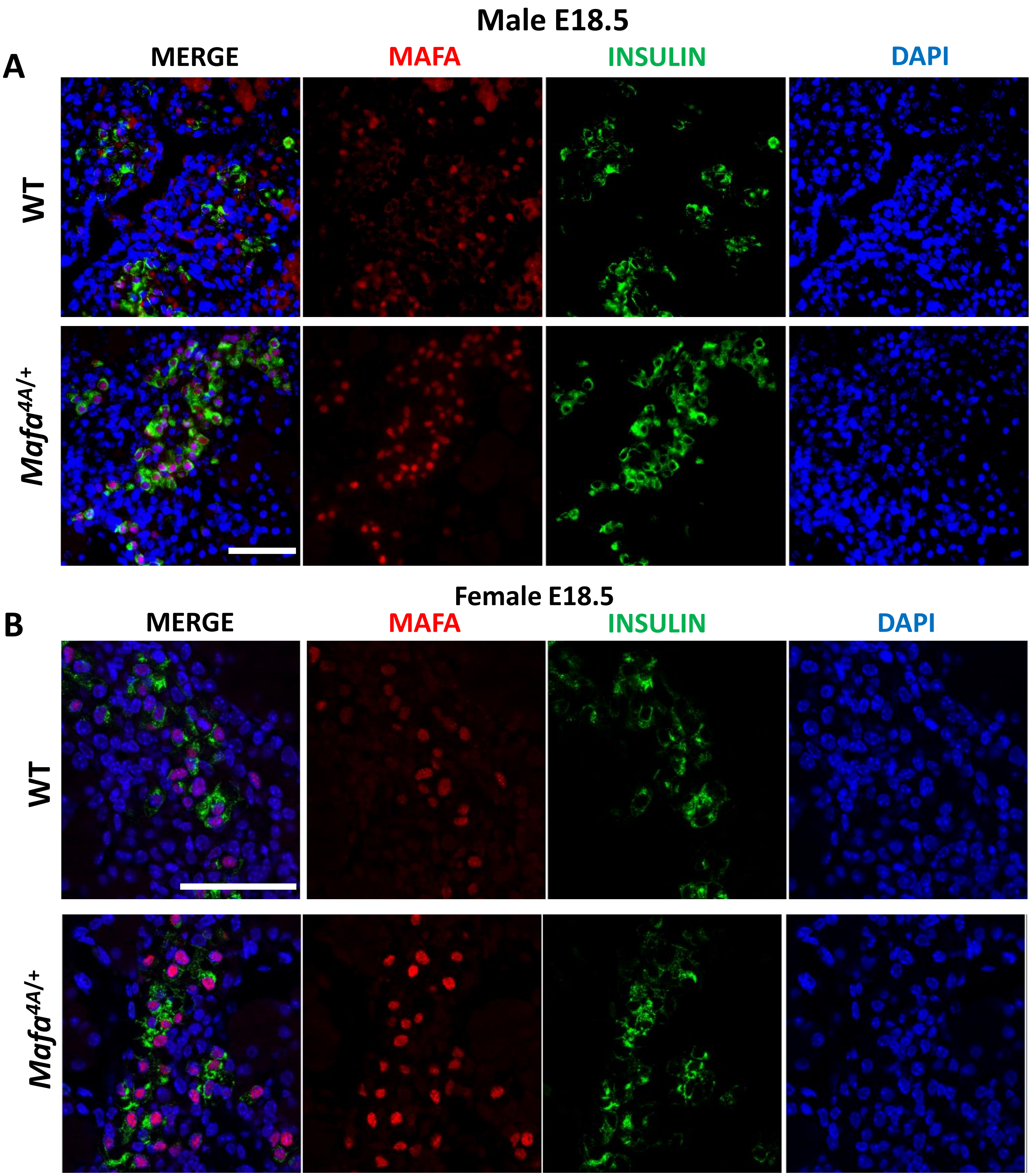

Fig SI 1 : Detection of MAFA in the foetal pancreas of males (A) and females (B) mice at stage E18.5. Immunofluorescence experiments were performed on pancreases isolated from *Mafa*$^{4A/+}$ mice and WT controls using anti-MAFA antibodies (red) and anti-insulin antibodies (green). Scale at 60µm

**Fig SI 2**

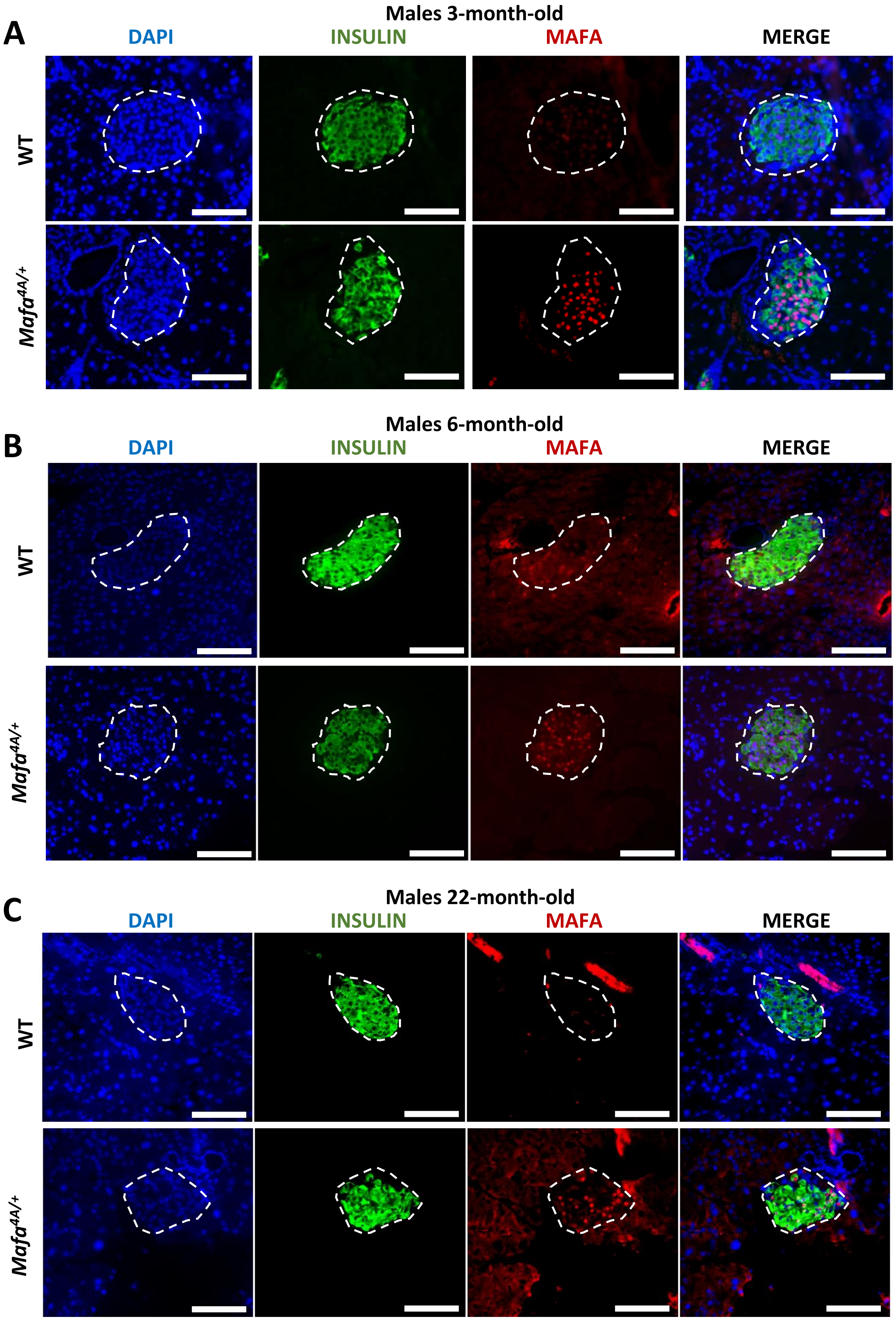

Fig SI 2 : Analysis of MAFA ontogenesis in male pancreases. Immunofluorescence staining was performed using anti-insulin antibodies in green and anti-MAFA antibodies in red at 3 (A), 6 (B) and 22 months (C) in WT and *Mafa*$^{4A/+}$ mice. Scale at 60µm

### Fig SI 3

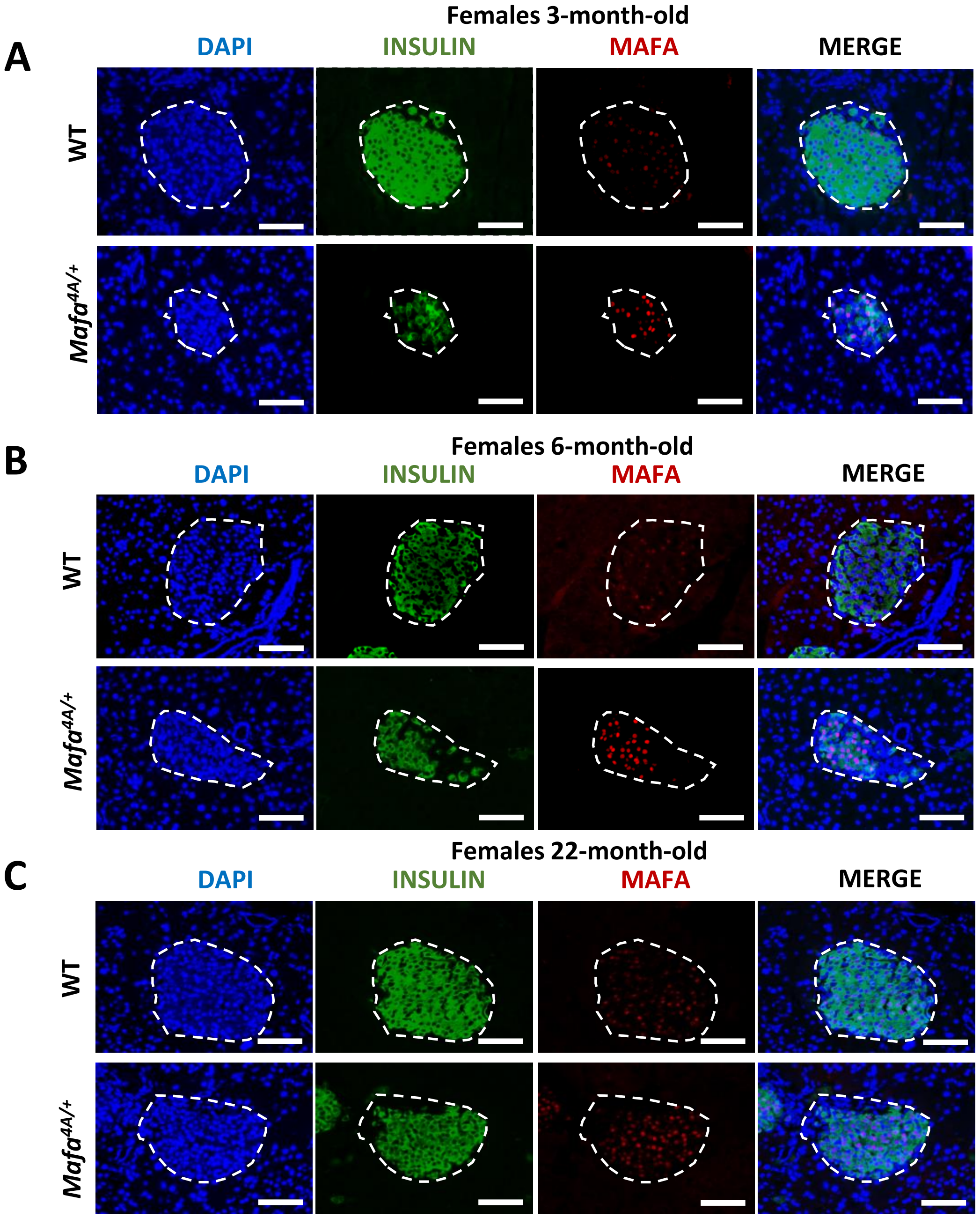

Fig SI 3 : Ontogenesis of MAFA in female pancreases. Immunofluorescence was performed on pancreases from *Mafa*$^{4A/+}$ and WT mice using anti-insulin antibodies in green and anti-MAFA antibodies in red at 3 (A), 6 (B) and 22 months (C). Scale at 60µm

### Fig SI 4

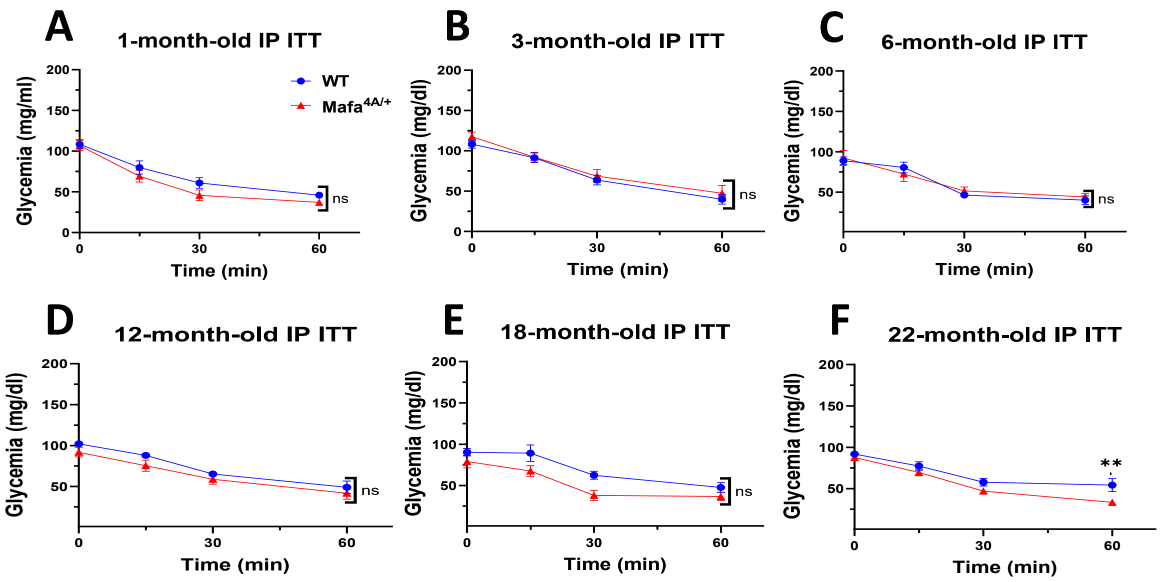

Fig SI 4: A-F) Intraperitoneal insulin tolerance test (IP ITT) performed after 6 hours of fasting on male mice at 1-month (A), 3-month (B), 6-months (C), 12-month (D), 18-month (E) and 22-month-old (F). Data represent a mean ± SEM of $n \geq 4$. 2-way ANOVA followed by Tukey's multiple-comparison test was used. **=$p<0.01$

### Fig SI 5

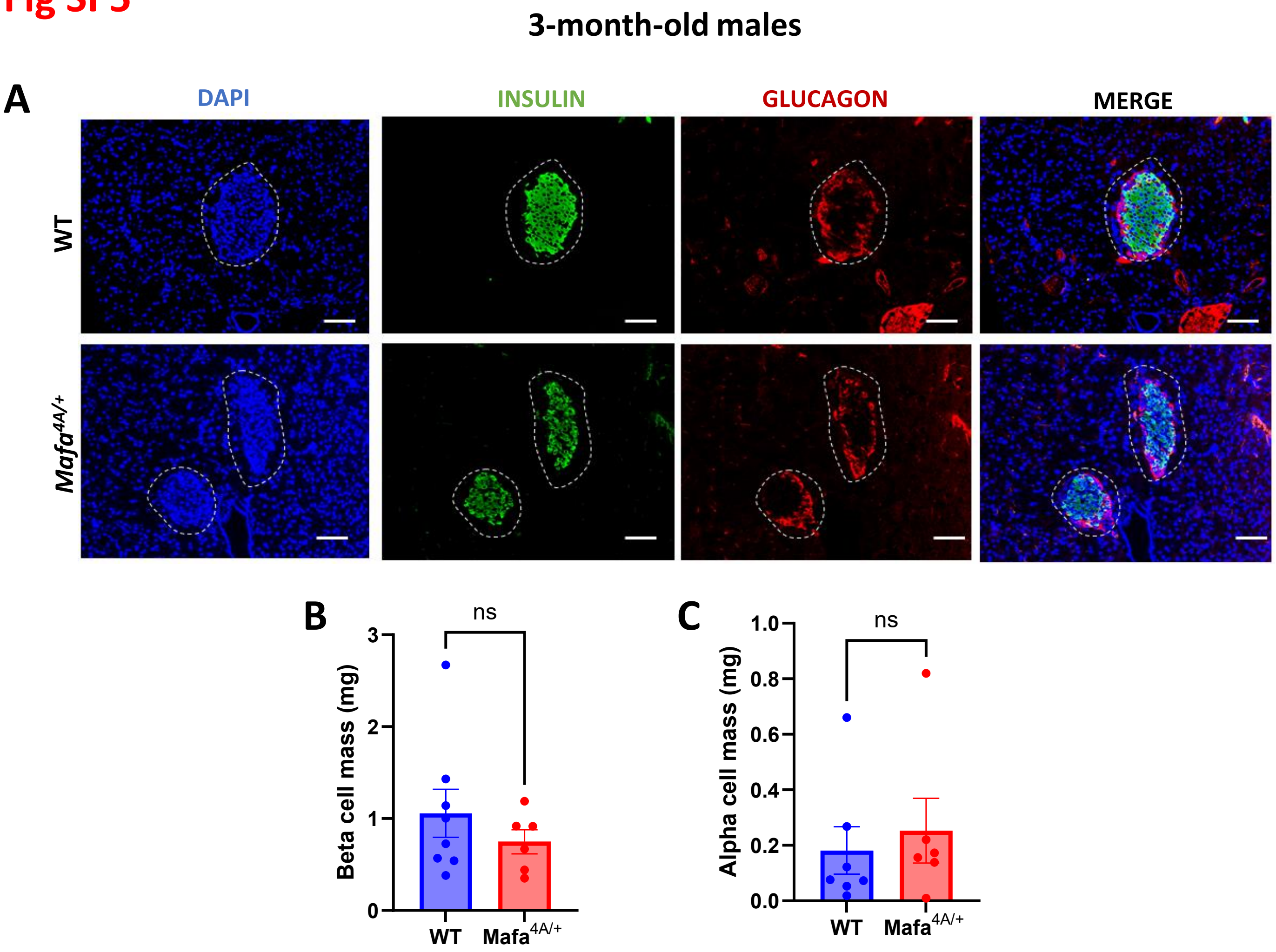

Fig SI 5 : Histological analysis of 3-month-old male mice pancreas. A) Immunostaining of beta cells with an anti-insulin antibody in green and alpha cells with an anti-glucagon antibody in red. B-C) Quantification of the beta-cells mass in B and alpha cell mass in C. Data represent a mean ± SEM of n >6. Unpaired t-test was used for C and Mann-Whitney test for B.

#### Fig SI 6

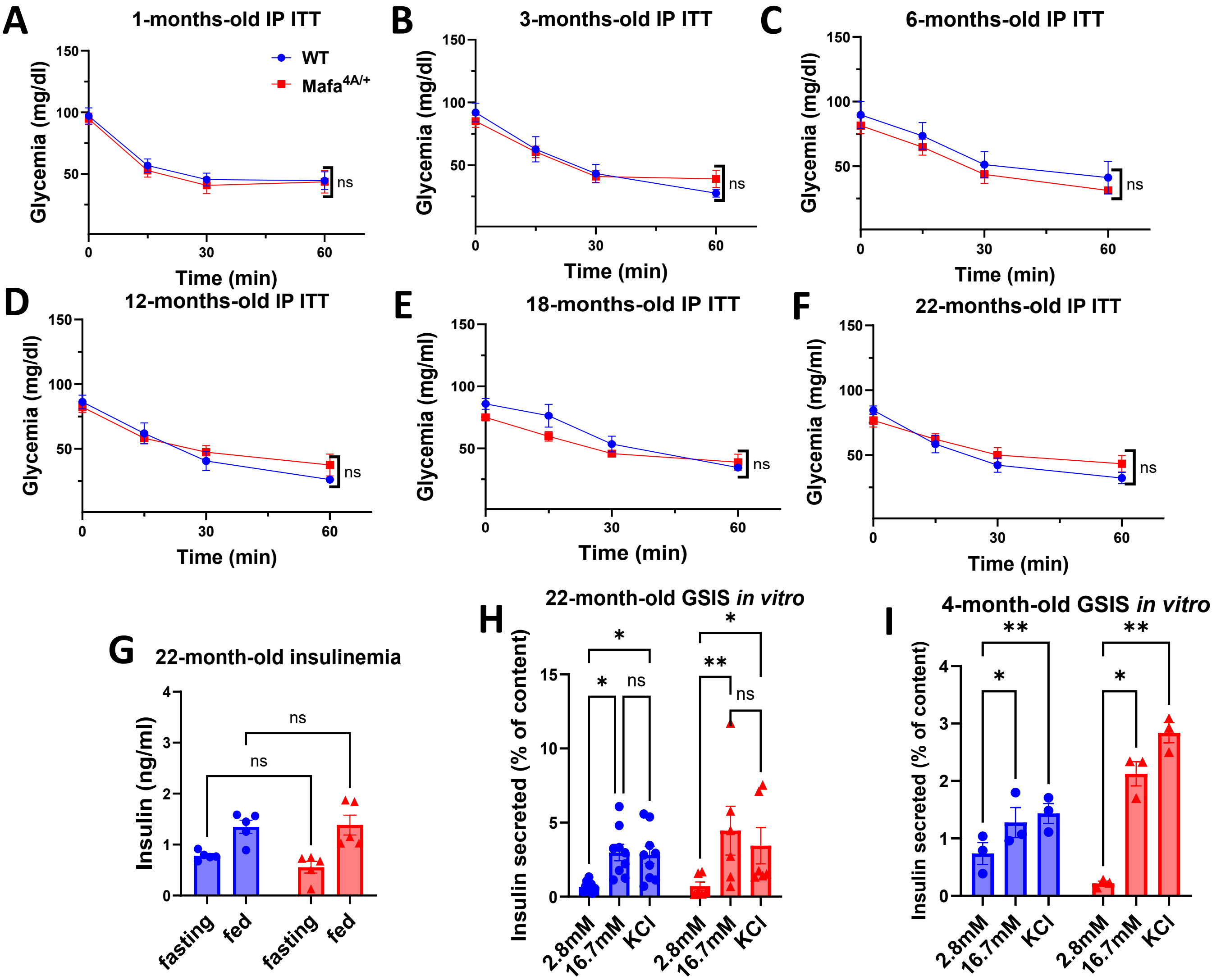

Fig SI 6 : Control of glucose homeostasis in females. A-F) Insulin tolerance test performed on female mice from 1 to 22 months of age. G) Insulinemia. H-I) GSIS performed on islets from WT and *Mafa*$^{4A/+}$ mice of 22-month-old (H) and 4-month-old (I). Data represent a mean ± SEM of n ≥ 3. 2-way ANOVA followed by Tukey's multiple-comparison test was used in A-I. *=p<0.05; **=p<0.01; ***=p<0.001; ****=p<0.0001.
